# Hyperactive mTOR in Lung Mesenchyme Induces Endothelial Dysfunction and Pulmonary Vascular Remodeling

**DOI:** 10.1101/2022.10.08.511300

**Authors:** Susan M. Lin, Ryan Rue, Akansha Goel, Kseniya Obraztsova, Alexander R. Mukhitov, Kanth Swaroop Vanka, Carly J. Smith, Maria C. Basil, Laura T. Ferguson, Joseph D. Planer, Edward Cantu, Grazyna Kwapiszewska, Edward E. Morrisey, Jillian F. Evans, Vera P. Krymskaya

## Abstract

Pulmonary vascular remodeling is the key structural abnormality in pulmonary hypertension (PH). Mechanistic target of rapamycin (mTOR) has long been suspected to play a role in the development of pulmonary vascular remodeling. However, underlying cellular and molecular mechanisms leading to this pathophysiological condition remain incompletely understood. To elucidate the crosstalk between lung mesenchyme with activated mTOR and endothelial cells (ECs), we focused on a monogenic lung disease, pulmonary lymphangioleiomyomatosis (LAM). LAM is a progressive cystic lung disease caused by a mutational inactivation of tuberous sclerosis complex (TSC1/TSC2), which results in constitutive mTOR activation in mesenchymal LAM cells. ECs derived from LAM lung explants showed increased proliferation, migration, and defective angiogenesis compared to age- and sex-matched ECs from control human lung. In LAM cells, we found increased WNT2 ligand expression. We also identified corresponding Frizzled 4 (FZD) receptors on ECs isolated from distal LAM lung, suggesting cellular crosstalk between LAM cells and ECs. In endothelial-fibroblast cocultures, treatment of normal ECs with WNT2 ligands recapitulated LAM EC phenotype and morphology. We observed transcriptomic upregulation in metabolic, angiogenic and growth pathways in ECs of young mice, while 1-year-old *Tsc2*^*KO*^ mice spontaneously developed pulmonary vascular remodeling with concurrent elevation in right ventricular systolic pressure. Our study demonstrates that LAM cells are not just a pathological mesenchymal cell state but a signaling hub that contributes to dysregulated cellular response in the surrounding vasculature, eventual pulmonary vascular remodeling and PH.

## Introduction

Pulmonary hypertension (PH) is a devastating lung disease characterized by pulmonary vascular remodeling^1,2^. Pulmonary vascular remodeling results in luminal narrowing, increased pulmonary vascular resistance, and ultimately progression to pulmonary hypertension and right heart failure^1-3^. Vascular remodeling occurs within the intimal and medial layers of lung vasculature through a complex and dynamic interplay between intimal ECs and medial smooth muscle cells^2^. Current PH treatments reduce pulmonary vascular resistance through vasodilation^4,5^, but none work to prevent or reverse pulmonary vascular remodeling. Development of new therapeutics remain limited as the molecular pathways regulating pulmonary ECs in acute and chronic lung diseases are not well defined. Recently, the mechanistic target of rapamycin (mTOR) pathway was implicated in the progression of pulmonary vascular remodeling in PH^6-9^, but how aberrant mTOR signaling leads to EC dysfunction and pulmonary vascular remodeling is not yet understood (in part due to heterogeneity in PH).

The mTOR pathway serves as a cellular sensor with increases in nutrients and growth-promoting factors such as amino acids, nucleotides, sugars, and insulin^10^. Mutational inactivation of the *TSC1/TSC2* gene, which is a key upstream negative regulator of mTOR activation^11,12^, is associated with the disorder tuberous sclerosis complex (TSC) and pulmonary lymphangioleiomyomatosis (LAM)^13-15^. LAM occurs predominantly in females and presents with cystic airspace enlargement, airflow obstruction and progressive lung function decline^16^. In lungs from patients with LAM, lesions formed by “LAM cells”, which harbor *TSC1* or *TSC2* mutations, consist of two cell subpopulations: myofibroblast-like spindle-shaped cells expressing smooth muscle-specific markers and epithelioid-like cells expressing glycoprotein gp100^17^. LAM cells are localized in lung parenchyma near cystic lesions, as well as pulmonary bronchioles, vasculature and lymphatics^18^. Approximately 10% of LAM patients develop pulmonary hypertension^18-20^ and histological studies have shown intimal and medial hypertrophy of pulmonary arteries^18^. However, there has been limited work to determine whether pulmonary vascular disease is intrinsically due to pathological mTOR activation in mesenchymal cells (WHO group 1 versus 5) or secondary to chronic lung disease (WHO group 3).

In this study, we investigated the effect of constitutive mTOR activation in lung mesenchyme on pulmonary ECs and lung function. We identified dysregulated WNT signaling as a potential driver for endothelial-mesenchymal crosstalk in primary distal vascular ECs isolated from LAM patients. Furthermore, we demonstrate that mTOR activation in the lung mesenchyme induces endothelial dysfunction in the small pulmonary vessels. To further interrogate the effects of lung-mesenchyme-specific mTOR hyperactivation on pulmonary ECs, we utilized a novel murine model of selective deletion of *Tsc2* in pulmonary mesenchymal progenitors (*Tbx4*^LME-Cre^*Tsc2*^fl/fl^)^21^. We previously found that *Tsc2*^*KO*^ mice with mesenchyme-specific loss of *Tsc2* developed age- and sex-dependent alveolar enlargement as well as other characteristic features of LAM lung disease including the emergence of a mesenchymal LAM cell with dysregulated mTOR activation, similar to human LAM cell subsets. In this study, we used this model to interrogate the effect of activated mTOR in pulmonary mesenchyme on pulmonary EC function. We demonstrated that mTOR activation in lung mesenchyme induces EC dysfunction through dysregulated WNT signaling to ECs, resulting in pulmonary vascular remodeling and pulmonary hypertension. Taken together, this suggests that mTOR hyperactivation directly causes EC dysfunction and the development and progression of pulmonary vascular remodeling and PH.

## Results

### ECs from LAM lung explants are characterized by a hyperproliferative phenotype with reduced angiogenic capacity

End-stage LAM lung explants contained remodeled vessels with intimal fibrosis and medial hypertrophy (Fig 1a). LAM nodules in the lung parenchyma were positive for pS6, a marker of mTOR activation^12^. To characterize ECs, peripheral LAM lung specimens with distal pulmonary vasculature were enzymatically dissociated into single cell suspensions as previously described^22-23^. ECs were isolated using magnetic bead sorts and grown to confluence on fibronectin-coated plates. Purity was confirmed by flow cytometry with canonical lineage markers and immunohistochemistry (Supplemental Fig 1). Compared to age- and sex-matched controls, LAM-derived ECs were more dysmorphic (Fig 1c) and proliferative (Fig 1d), with increased migration (Fig 1e-f). Next, the angiogenic capacity of ECs was evaluated using an *in vitro* tube formation assay (Fig 1g). On these 2-dimensional monocultures, LAM ECs exhibited decreased tube lengths compared to control ECs, indicating reduced angiogenic potential when grown in the absence of support cells (Fig 1h).

**Figure 1.**
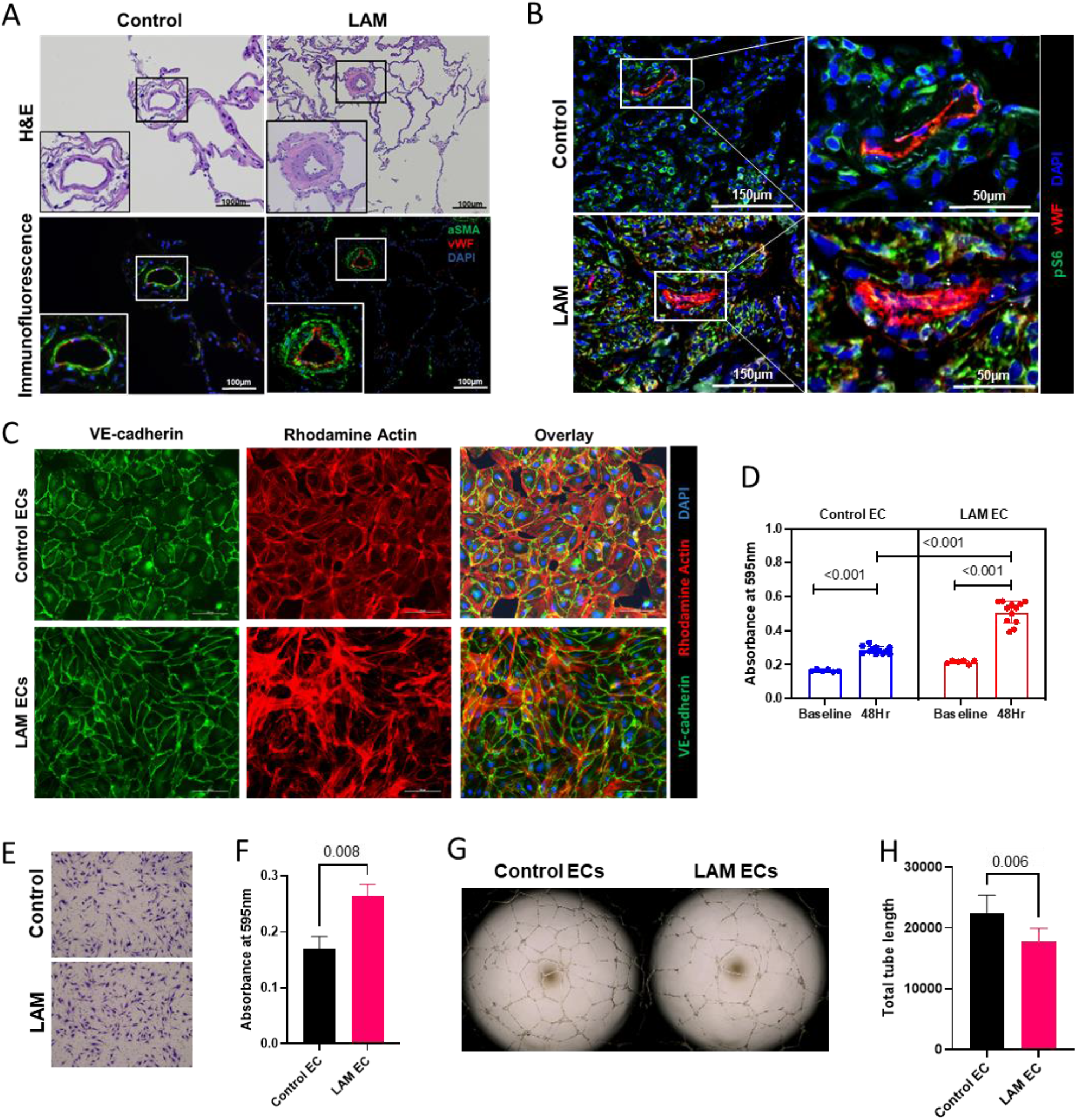
Characterization of pulmonary ECs from patients with LAM. (a) Immunohistochemistry (IHC) of distal vasculature notable for increased intimal fibrosis and medial hypertrophy in LAM lung compared to age- and sex-matched control lung. Immunofluorescence staining for ECs (von Willebrand Factor; red) and smooth muscle (α-smooth muscle actin; green). (b) IHC with pS6 (marker of mTOR activation) and vWF (endothelial marker). (c) LAM and control pulmonary ECs immune-labelled with antibodies against endothelial marker vWF (green) and F-actin (rhodamine phalloidin; red). Nuclei countered using DAPI (blue). (d) Proliferative capacity of normal and LAM ECs over 24 hours using a crystal violet growth assay. (e) Representative images of migration assay of normal and LAM EC performed on Boyden chambers using inserts with 0.8µm pores. (f) Migration assays. After 24 hours, migrated cells were fixed with 2.5% glutaldehyde and stained with 0.05% crystal violet with absorbance subsequently read at 595nm. (g) Representative image of *in vitro* angiogenic potential of LAM and control ECs using angiogenesis assay. (h) Total tube length analyzed using Angiogenesis Analyzer plugin on Image J.

### LAM lung fibroblasts increase angiogenic potential of LAM ECs

To assess endothelial-mesenchymal crosstalk through cell-cell and cell-matrix interactions, primary ECs and fibroblasts were co-cultured on fibronectin-coated 12-well plates (Fig 2a). In coculture, LAM ECs demonstrated increased affinity for self-segregation and self-organization (Fig 2b). LAM ECs had greater cell counts than control ECs when cocultured with control fibroblasts, which increased further when LAM ECs were cocultured with LAM fibroblasts (Fig 2c). ECs within LAM EC and fibroblast cocultures occupied more cell surface compared to control ECs in coculture with control fibroblasts (Fig 2d). The ability of LAM ECs to self-assemble and form endothelial tubes was then assessed within 3-dimensional (3D) extracellular matrix (ECM)-based scaffolds (Fig 2e). When grown in isolation, there were no significant differences in total tube length and cell count between control and LAM ECs (Fig 2f). However, when grown in cocultures with fibroblasts, LAM ECs developed more extensive tube networks (Fig 2g). Taken together, the major changes in EC organization in 2D cocultures and the significant increase in cell count and tube formation in 3D cocultures demonstrate that mesenchymal cells from LAM lung induce EC proliferation and *in vitro* vasculogenesis.

**Figure 2.**
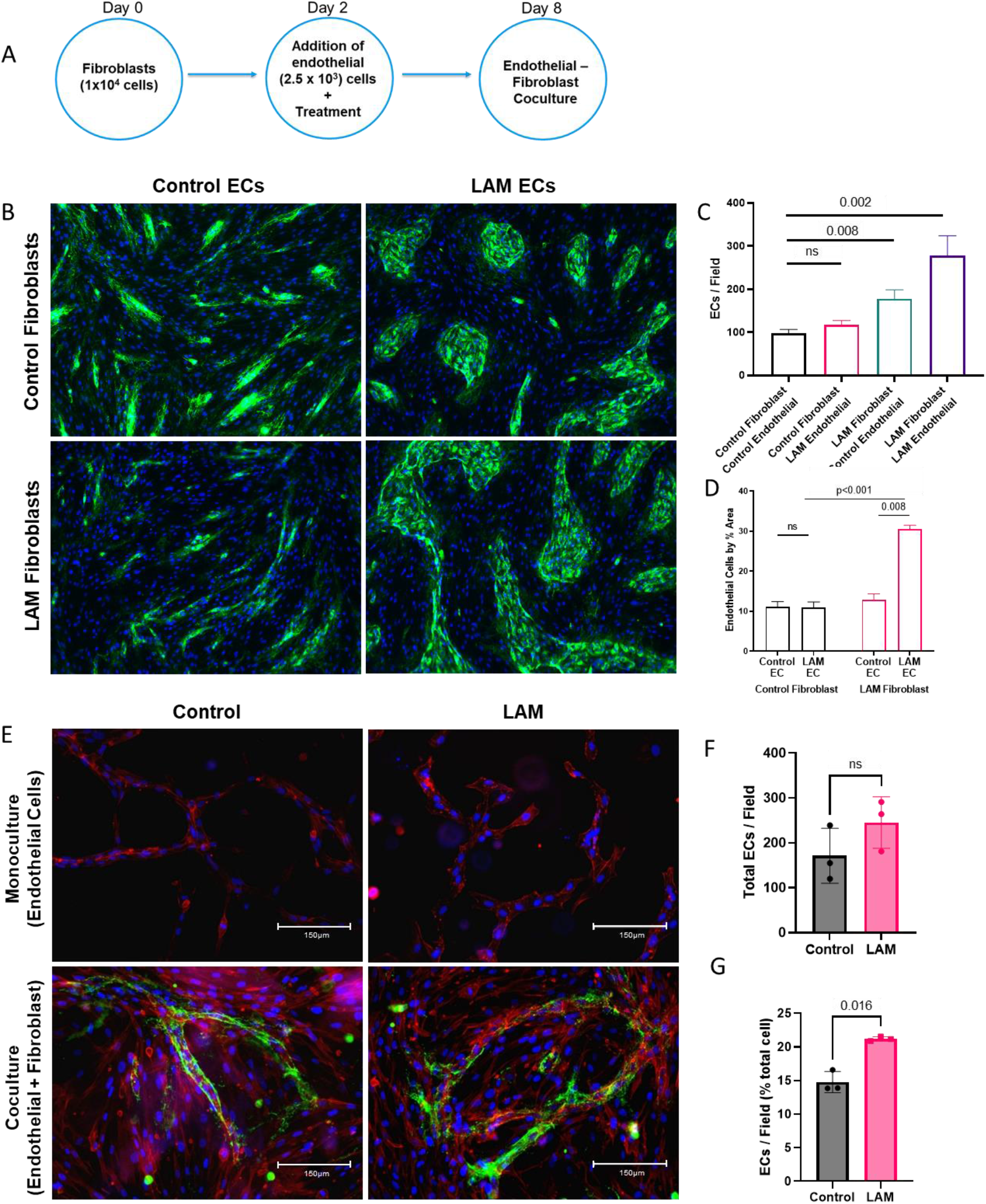
Increased EC proliferation and *in vitro* vasculogenesis in three-dimensional (3D) endothelial-fibroblast coculture systems. (a) Schematic for 2D endothelial-fibroblast coculture; where 9×10^4^ fibroblasts were plated and allowed to settle prior to addition of 3×10^4^ ECs; then cocultured for 8 days. (b) Representative images of endothelial-fibroblast cocultures using primary cells isolated from LAM lung explants as well as age-matched control lungs. ECs stained with vWF (green) and nuclei counterstained with DAPI (blue). (c) Number of ECs/field in the endothelial-fibroblast cocultures. LAM ECs were more proliferative when cocultured with LAM fibroblasts (p=0.002). This effect was also seen when normal ECs were cocultured with LAM fibroblasts (p=0.0087). (d) EC by surface area in coculture conditions. % area measured on Image J. (e) Representative images of ECs grown with and without fibroblasts to create 3D sphericals in serum-free gel matrix. ECs per field in 3D sphericals with greater cell counts in LAM sphericals compared to normal sphericals. EC stained with vWF (green) (f) Total number of ECs per field (n=3) in normal compared to LAM 3D sphericals. (g) ECs/field (% of total cells) in 3D cocultures. LAM ECs were more proliferative when grown in the presence of LAM fibroblasts. Statistical analysis was performed using the nonparametric Kruskal-Wallis ANOVA test with Siegel (Bonferroni) correction for post-hoc, pair-wise contrasts n=3 control, 3 LAM ECs.

### LAM scRNA-seq reveals transcriptomic alterations in pulmonary vascular ECs

We analyzed available LAM single cell RNA sequencing data^21^ for transcriptomic changes in mesenchymal cell and EC clusters. The four major cellular compartments, specifically epithelial, immune, mesenchymal, and endothelial cells, were identified using canonical markers^26-28^. Endothelial cells were subsequently extracted and clustered for further analysis based on cell-selective markers^27-29^. In brief, pulmonary arterial endothelial cells were characterized by high expression of *GJA5* and *DKK2* genes; venous ECs by *ACKR1* and *HDAC9*; lymphatic ECs by *PROX1*, capillary (CAP) ECs by *ILJR* (CAP1) or *EDNRB* (CAP2); and Car4-high^30^ (Car4^+^) ECs by *CA4* and *CD34*. In this analysis, we found increased WNT2 ligand expression in LAM cells as well as their corresponding Frizzled 4 (FZD) receptors on ECs in LAM specimens (Fig 3a and Supplemental Fig 2). WNT signaling pathways have a key role in the preservation of pulmonary vascular homeostasis^31^. The presence of WNT activation was confirmed by *in-situ* hybridization with RNAscope of the WNT pathway marker AXIN2 (Supplemental Fig 2). WNT target gene Wnt2 (Fig 3b) was significantly increased in LAM lung mesenchyme compared to normal lung mesenchyme (Fig 3c).

**Figure 3.**
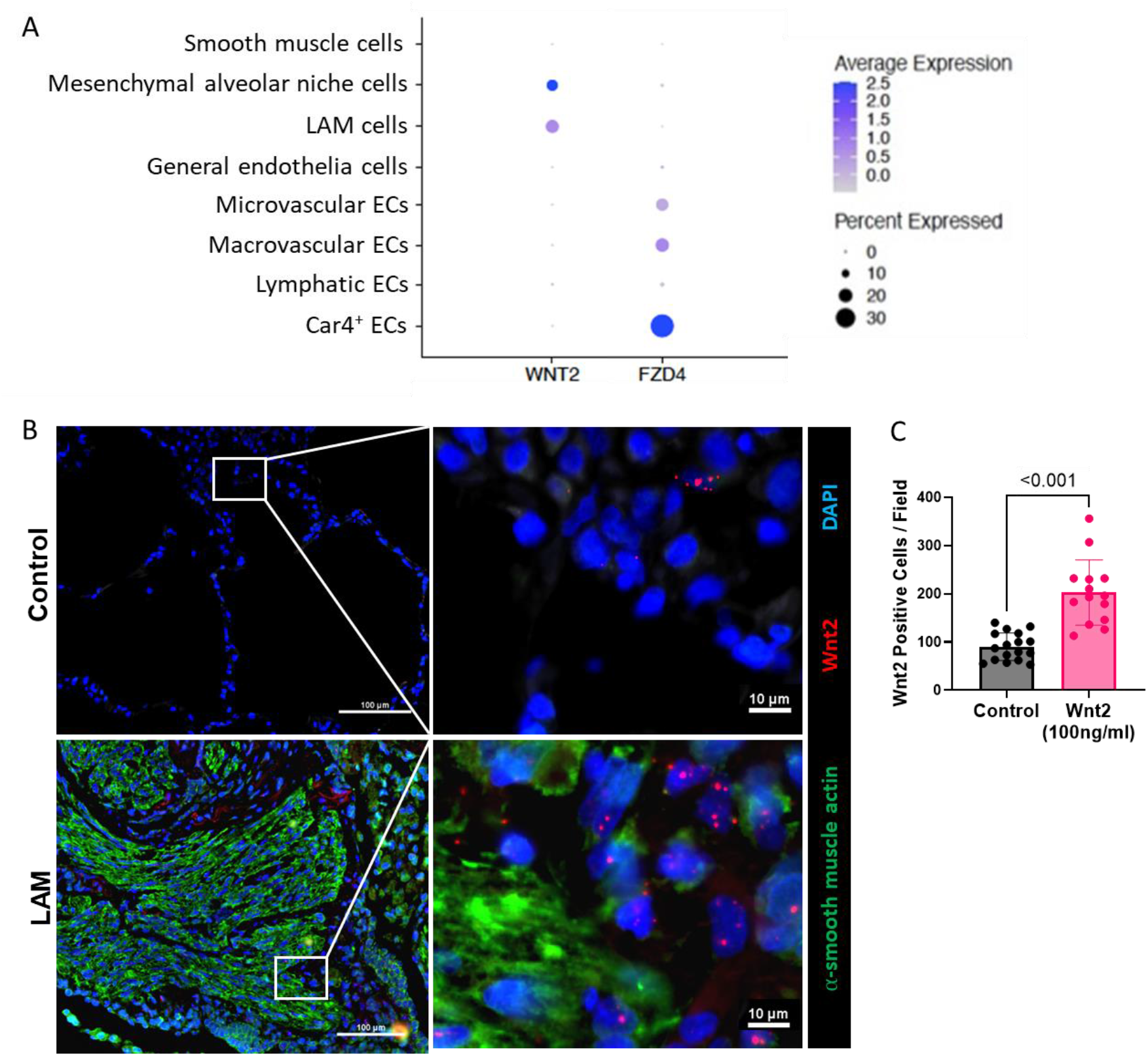
WNT pathway activation in LAM lung mesenchyme. (a) Wnt2 activation in LAM cell cluster within LAM lungs. Increased expression of Wnt2 in LAM cells with corresponding increased Fzd4 (receptor) expression in endothelial cells (b) WNT2 in Situ Hybridization of LAM and control human lung with increased expression within LAM lesions. (c) Percent of cells with WNT2 positivity in control compared to LAM lungs.

### WNT2 treatment of control ECs recapitulated LAM EC morphology

To assess the effect of mTOR and WNT pathway inhibition *in vitro* on EC function, LAM ECs were treated with C82, a WNT pathway transcriptional inhibitor, (Fig 4a) which decreased cell proliferation (Fig 4b), migration (Fig 4c-d), and endothelial tube formation (Fig 4e-f). LAM EC cocultures were more susceptible to mTOR inhibition than control EC cocultures, but there was no significant difference in response to WNT inhibition (Fig 4g). When cocultures with control ECs and control fibroblasts were treated with WNT2 ligand (100ng/ml), there was an increase in number of ECs (Fig 4g-h). In addition, treatment of control endothelial-fibroblast cocultures with WNT2 recapitulated LAM EC phenotype and morphology (see Fig 2b). Taken together, these experiments highlight the ability of control human ECs to adopt LAM EC morphology when activated by WNT2.

**Figure 4.**
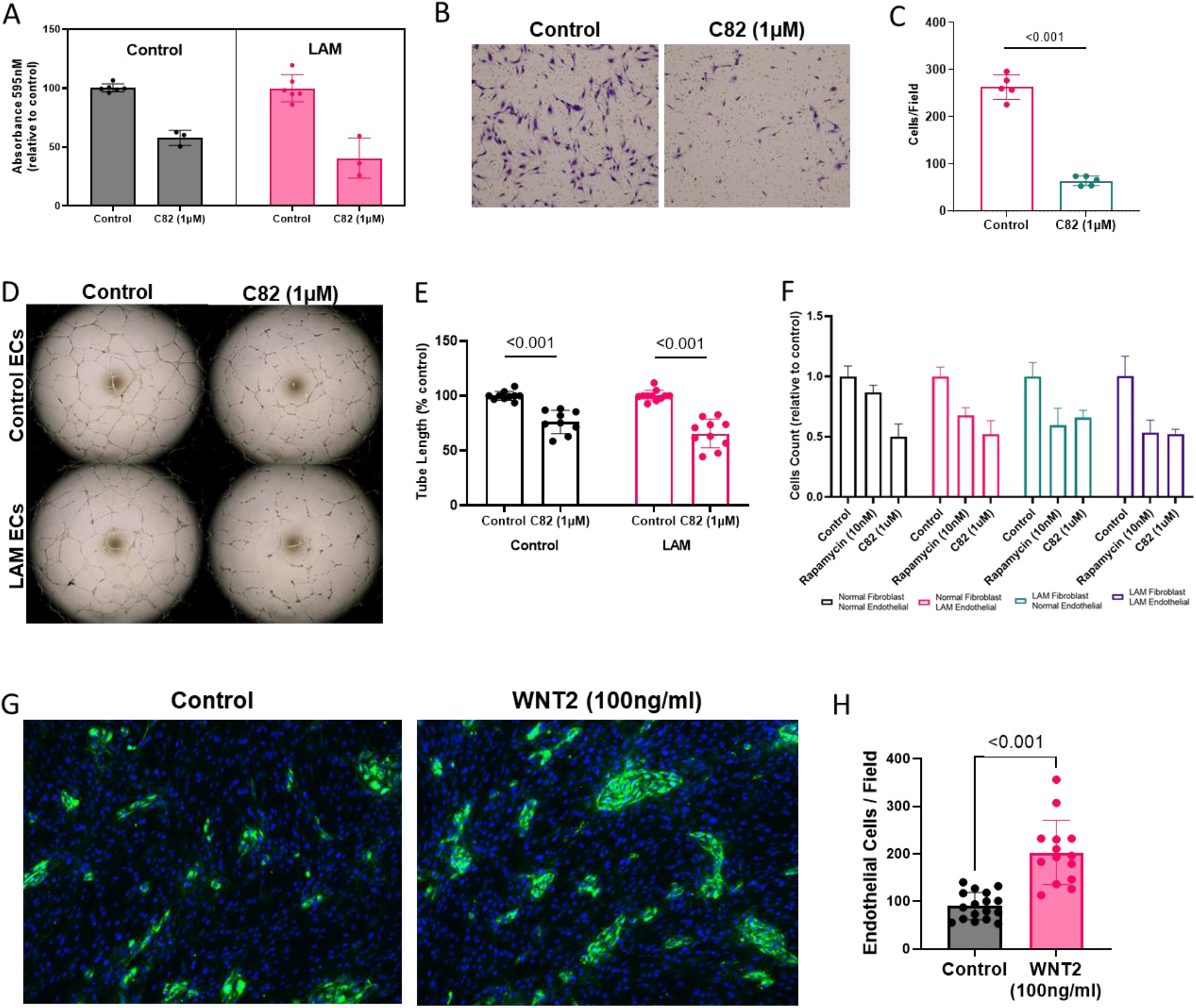
Inhibition of WNT suppresses EC proliferation, migration and angiogenesis. (a) Proliferation assay after treatment with WNT inhibitor C82 (1uM). (b) Representative images of migration assay with 1uM C82 inhibition. (c) Migration assay with cells/field. (d) Representative images from angiogenesis assay of normal and LAM ECs treated with 1uM C82. (e) Inhibition of WNT signaling with C82 decreases EC tube formation on 2D angiogenesis assay. Analysis of total tube length performed using Angiogenesis Analyzer plugin on Image J. (f) Decreased cell counts on endothelial-fibroblast cocultures when treated with 10nM rapamycin and 1μM C82. (g) LAM coculture phenotypes was recapitulated in normal endothelial-fibroblast cocultures when treated with WNT2 (100ng/ml). (h) Increased ECs per field on cocultures when treated with WNT2.

### Upregulation of metabolic, growth and angiogenic pathways in ECs from 8-week-old *Tbx4Tsc2*^*KO*^ mouse lung

To determine the mechanisms driving endothelial dysfunction in mTOR-activated lungs, we utilized *Tbx4*^LME-Cre^*Tsc2*^fl/fl^ mice^21^, a novel model of mTOR hyperactivation with targeted *Tsc2* deletion in the lung mesenchymal progenitor niche. mTOR activation was confirmed by presence of pS6 in *Tbx4*^LME-Cre^*Tsc2*^fl/fl^ (*Tsc2*^KO^) but not in T*bx4*^LME-Cre^*Tsc2*^WT/WT^ (*Tsc2*^WT^) mice (Fig 5a). We performed bulk RNA sequencing to characterize potential transcriptomic alterations in EC subpopulations. Pulmonary ECs from *Tsc2*^*KO*^ mice showed marked upregulation in metabolic, growth and angiogenic pathways compared to lung ECs from *Tsc2*^*WT*^ mice (Fig. 5b). Analysis of the top differentially expressed genes in *Tsc2*^*KO*^ pulmonary ECs exhibited upregulation of WNT ligands and amplifiers including Axin2, LRP6 and the frizzled WNT receptors Fzd1, Fzd2, Fzd7 and Fzd8 (Fig 5c).

**Figure 5.**
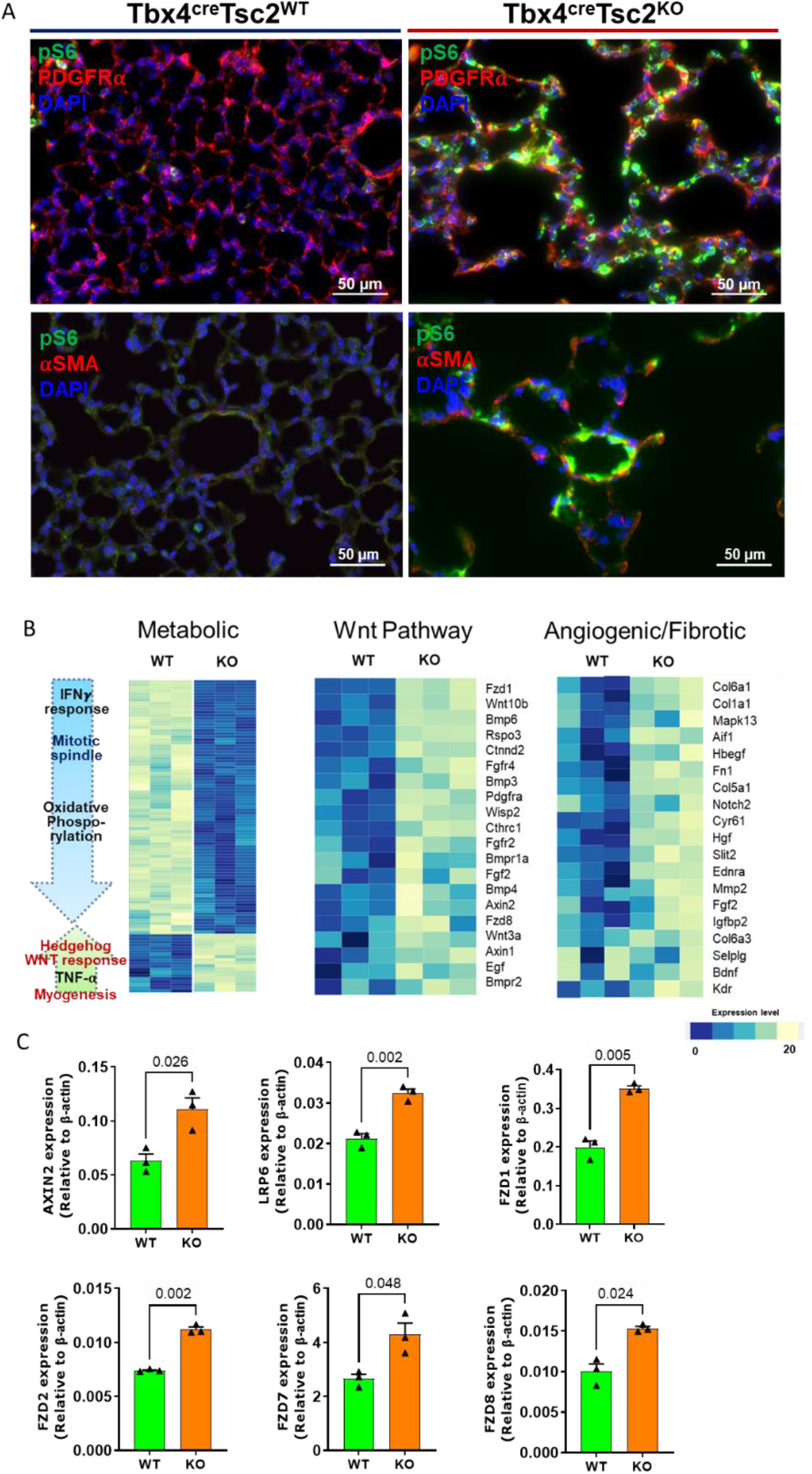
Differential gene expression changes in the CD31+ vascular ECs. (a) Representative immunohistochemical images of alveolar changes in lung mesenchymal-specific mTORC1 activation. Lung analysis demonstrates colocalization of mTORC1 activation (pS6) in PDGFRβ positive cells in *Tsc2*^KO^ mice. (b) Heatmap of the top differentially expressed genes in the mouse lung ECs isolated from wildtype (n=3) and knockout (n=3) mouse lungs. Upregulation of genes involved in metabolic, growth and angiogenic pathway in ECs from *Tsc2^KO^* mice. (c) RT-PCR of Wnt ligands in 8-week-old female mice.

### Targeted mesenchymal *Tsc2* deletion altered EC angiogenesis but had no significant effect on vessel number or peripheral muscularization at 12-, 16- and 20-weeks of age

In primary pulmonary ECs derived from 8-week-old mouse lungs, *Tsc2*^KO^ ECs form fewer capillary-like structures and tubes (Fig 6a). *Tsc2*^WT^ ECs form tubes within 8 hours while *Tsc2*^KO^ tube formation was not complete until 18 hours (Fig 6b). Tube formation was dependent on WNT signaling, with tube formation inhibited by C82 (Fig 6c). Vascular muscularization was scored based on the ratio of smooth muscle around vascular diameter (Fig 6d). Although loss of alveolar surface area can cause vascular pruning, there were no significant differences in the number of vessels or proportion of fully muscularized vessels in *Tsc2*^KO^ at 12-, 16- and 20-weeks of age (Fig 6e-f). These results suggest that while there were no histopathological changes at younger ages, isolated ECs from young *Tsc2*^KO^ mice were functionally altered by the exposure to constitutively activated mTOR in pulmonary mesenchyme.

**Figure 6.**
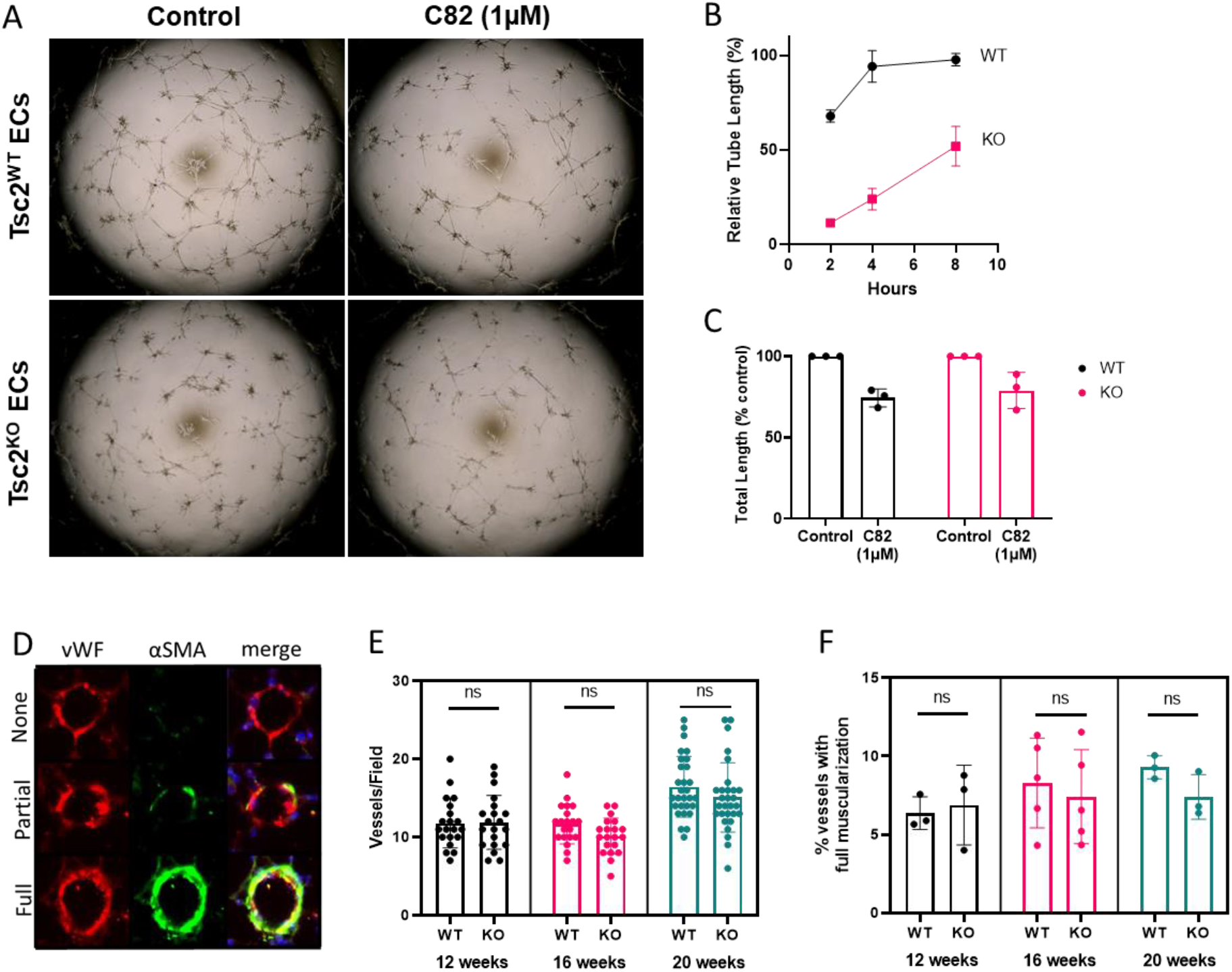
Lung mesenchymal-specific mTORC1 activation did not alter vasculature count or remodeling at 8weeks but significantly altered EC function. (a) Representative images of angiogenesis tube formation with ECs treated with 1uM C82. (b) Change in total tube length following treatment with C82. (c) Time to maximal tube formation (n=3) in control WT versus KO mice. (d) Representative figures of scoring metric for vascular remodeling based on smooth muscle involvement. The degree of muscularization was defined by α-smooth muscle actin positive parts as percentage of the total pulmonary artery cross section: non-muscularized: < 20%, partial muscularization: 20-70%, fully muscularized: > 70%. (e) There were no significant difference in total number of vessels or (f) vascular muscularization.

### Mesenchymal mTOR activation in lungs of one-year-old *Tsc2*^*KO*^ mice leads to spontaneous pulmonary vascular remodeling, elevated right ventricular systolic pressure and right ventricular hypertrophy

Although a marked alteration in transcriptomic signatures did not translate to significant vascular pathology up to 20 weeks of age in *Tsc2*^*KO*^ mice, we hypothesized that single cell and bulk RNAseq profiles may reflect early endothelial changes prior to histopathological changes at the end-stage disease. Analyses of 54-week-old *Tsc2*^*KO*^ mouse lungs (Fig 7a) demonstrate significant vascular remodeling compared to age-matched *Tsc2*^*WT*^ controls (Fig 7b-c). This was characterized by an increase in fully muscularized vessels with subsequent decrease of non-muscularized vessels, confirmed using manual counts of selective immunohistochemical staining of vessels (Fig 7b) and automated analyses using Visiomorph (Fig 7c). In addition, pulmonary vessels in *Tsc2*^*KO*^ mice had thicker medial walls (Fig 7d). However, there were no differences in the total number of vessels (Fig 7e). We investigated right ventricular systolic pressures in 54-week-old mice by right heart catheterization and found elevated ventricular pressures in *Tsc2*^*KO*^ compared to *Tsc2*^*WT*^ (Fig 7f). *Tsc2*^*KO*^ also had larger hearts measured by gross weight (Fig 7g), with higher right ventricular mass as measured by Fulton Index (Fig 7h). Fulton Index, a metric of right ventricular hypertrophy, was most notable in female *Tsc2*^*KO*^ mice (Fig 7i).

**Figure 7.**
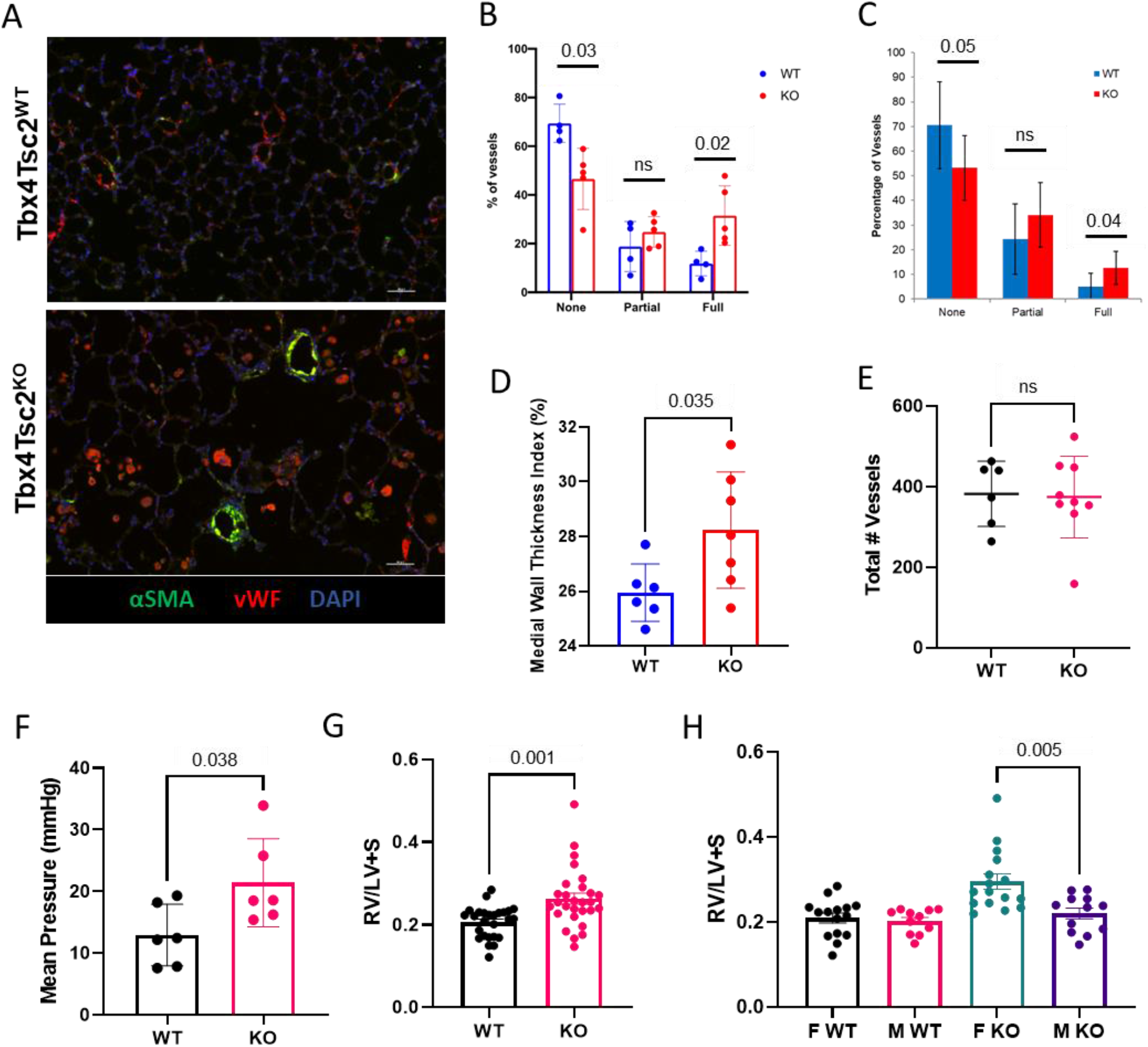
Pulmonary vascular remodeling and right heart dysfunction in one-year-old Tbx4Tsc2^KO^ mice with mesenchymal mTOR activation. (a) Representative images of vessels in *Tsc2^WT^* compared to *Tsc2^KO^* in one-year-old mice. (b) Scoring of peripheral muscularization based on 15 randomly acquired images per mouse (n=9). (c) Peripheral muscularization based on VisiomorphTM software (Visiopharm; Hoersholm, Denmark). Morphometric analysis of pulmonary vascular remodeling in *Tsc2^KO^* (n=9, red) mice compared to *Tsc2^WT^* mice (n=6, blue) mice. (d) Medial wall thickness based on Visiomorph. Medial wall thickness was determined in vessels with a diameter of 20–70 μm as well as the lumen area (defined as the area within the lamina elastic interna). Comparison of *Tsc2^WT^* (n=6) to *Tsc2^KO^* n=7). (e) Total number of vessels unchanged in *Tsc2^WT^* vs *Tsc2^KO^*. (f) Right heart catherization of one-year-old mice (6 *Tsc2^WT^*, 6 *Tsc2^KO^*) with measurements of right ventricular systolic pressure. (g) Right ventricular hypertrophy (RVH) as measured by Fulton index (RV/S+LV); *Tsc2^WT^* (n=25) vs *Tsc2^KO^* (n=27). (h) Increased Fulton index were driven by females *Tsc2^KO^* (n=13) compared to male *Tsc2^KO^* (n=11).

## Discussion

Management of pulmonary hypertension has long been limited by the inability of current therapies to prevent or slow progression of vascular remodeling^32^. Using human samples and a murine model, our study is the first to explore the effects of concomitant mTOR and WNT signaling on the crosstalk between mesenchymal and pulmonary endothelial cells in the development of pulmonary vascular remodeling. In the presence of hyperactivate mesenchymal mTOR, there is increased proliferation and migration of LAM ECs compared to control ECs, but reduced angiogenic capacity in monoculture angiogenesis assay. The combination of hyperproliferation with reduced angiogenic capacity has previously been observed in primary pulmonary endothelial cells isolated from patients with idiopathic pulmonary hypertension^24^ and chronic thromboembolic pulmonary hypertension^25^. To study biochemical and biophysical cues that promote vascularization, we developed (1) endothelial-fibroblast cocultures to study cell-cell interactions and (2) endothelial-fibroblast cocultures in 3D hydrogels to study cell-matrix interactions on a microvascularized model. We found that when cocultured with fibroblasts, LAM ECs had significantly enhanced growth and drastically altered cellular segregation. This effect was further augmented in the 3D matrix model with increased EC tube length in LAM ECs grown with LAM fibroblasts.

To understand the implications of endothelial dysfunction in a mTOR hyperactivated mesenchymal state, we turned to a murine model of selective mTOR activation in mesenchymal cells. In these *Tsc2*^*KO*^ mice, we found that transcriptomic upregulation of metabolic, growth factor and angiogenic genes occurred by 8 weeks of age; however, these mice do not exhibit vascular muscularization until 1 year of age. Intriguingly, this is comparable to a human age of ∼40-50 years^33-34^, the average age of respiratory failure in patients with LAM requiring initiation of allosteric mTOR inhibitor sirolimus^35^. Thus, our murine studies confirmed the ability of mTOR hyperactive mesenchymal-derived fibroblasts to alter transcription and function in ECs, resulting in spontaneous pulmonary vascular muscularization and PH in the lungs of 1-year-old mice.

The presence of transcriptomic changes prior to onset of pulmonary vascular remodeling offered a unique opportunity to study early disease state. We identified the WNT ligand, Wnt2, as a potential activator in LAM mesenchyme. The WNT/planar cell polarity pathway is known to determine the vasculogenic fate of post-natal mesenchymal stem cells^36^ with proper endothelial cell differentiation^37^ and angiogenesis^38^ contingent on WNT/β-catenin signaling. Correspondingly, in human EC studies, we demonstrated that WNT inhibition limited LAM EC proliferation, migration and angiogenesis. The apparent crosstalk between ECs and fibroblasts in cocultures was inhibited by blocking either WNT or mTOR signaling. Lastly, WNT2 activation elicited a LAM-like phenotype *in vitro*, characterized by increased cell count and alterations in cellular morphology. Taken together, these findings confirmed the importance of the WNT pathway in EC function and demonstrated that activation of the WNT/β-catenin pathway recapitulated LAM EC dysfunction.

Our work highlights the role of both mTOR and WNT2 in the progression of vascular remodeling in LAM and hyperactive mTOR lung diseases, but further work is needed to extrapolate these findings to other diseases that cause pulmonary hypertension. Specifically, pulmonary ECs are a heterogeneous collection of cells and as such, require additional research into individual population responses to physiological and pathological stimuli. Moreover, while our study focused on endothelial cells, pulmonary artery smooth muscle cells (PASMC) have an equally important structural and functional role in the development and progression of pulmonary vascular remodeling. Recent research on pulmonary vascular remodeling suggest a phenotype shift in PASMCs from contractile to synthetic phenotypes.^39^ Further work is needed to understand EC and PASMC heterogeneity and how it contributes to vascular homeostasis in health and disease states.

These studies demonstrate that LAM ECs are dysfunctional despite constitutive mTOR activation occurring in only a subset of lung mesenchymal cells. Next generation sequencing allowed us to characterize mechanistic pathways in an agnostic fashion, leading to the discovery of WNT/β-catenin and specifically, WNT2, as an activator of EC dysfunction. We subsequently utilized a murine model of mesenchymal mTOR activation to demonstrate endothelial dysfunction as well as spontaneous pulmonary vascular remodeling. Taken together, these studies demonstrate that LAM cells, a pathological cell subset with constitutive mTOR hyperactivation, drive cellular dysfunction in other cell subsets. Through dysregulated paracrine signaling from LAM cells to ECs, LAM cells act through WNT pathway to contribute to pulmonary vascular remodeling and PH. This highlights that LAM cells are both a pathological mesenchymal cell state observed in disease and a signaling hub that promotes dysregulated cellular response in the surrounding parenchymal and contributes to eventual pulmonary vascular remodeling and PH.

## Methods

### Experimental animals

All animal procedures were approved by the University of Pennsylvania Institutional Animal Care and Use Committee and conform to NIH guidelines on animal care. As previously described^21^, *Tsc2*^*loxP/loxP*^ mice were crossed with *Tbx4*^*LME_Cre*^ mice. *Tsc2*^*KO*^ referred to *Tbx4*^*LME_Cre*^ *Tsc2*^*loxP/loxP*^ mice while *Tsc2*^*WT*^ mice referred to *Tbx4*^*LME_Cre*^ *Tsc2*^*WT/WT*^ mice.

### Mouse lung histology and morphometry preparation

Lungs were inflated with under constant pressure (25 cm H2O) and the pulmonary circulation was flushed with 2 cc of PBS as described^21,40^.

### Human lung tissue

Normal lung samples were obtained through the University of Pennsylvania Lung Biology Institute’s Human Lung Tissue Bank. These samples were obtained from deceased donors and represent secondary use of tissue. Lymphangioleiomyomatosis (LAM) lung samples were obtained from living donors at the time of lung transplantation through the National Disease Research Institute (NDRI, Philadelphia Pennsylvania). Informed consent was obtained by NDRI prior to acceptance of tissue donation for research. For both normal and diseased (LAM) lung samples, identifying information were removed prior to use in accordance with institutional and NIH protocols.

### Tissue and scRNA-seq sample preparation

Lung parenchymal tissue preparation and scRNA-seq sample preparation were performed as previously described.^22-23^ In brief, human lungs specimens were first confirmed to be from the most distal portion of the lung through direct visualization of the pleural lining. The pleural lining was separated from the lung parenchyma and discarded followed by microdissection along the bronchovascular tree. Specimens were then mechanically minced prior to enzymatic digestion with collagenase, dispase and DNase. Samples were then filtered, RBC lysed with ACK lysis buffer and processed into single cell suspension. CD45+ immune cells were depleted using MACS LS columns with CD45-microbeads (Miltenyi, 130-045-801) with 2 × 10^6^ cells per column to enhance purity and viability. Following single cell suspension, the samples were loaded onto a GemCode instrument (10x Genomics) to generate single-cell barcoded droplets (GEMs) according to the manufacture’s protocol. The resulting libraries were sequenced on an Illumina HiSeq2500 or NovaSeq instrument. Reads were aligned, and gene-level unique molecular identifier (UMI) counts were obtained using the Cell Ranger pipeline. Analyses were performed using the R package Seurat v3.

### Immunohistochemistry

Human and murine paraffin-embedded tissue sections used for the immunohistochemistry experiments were sectioned and deparaffinized in xylene followed by isopropanol dilutions. Antigen retrieval was performed with 10 mM sodium citrate buffer (10 mM sodium citrate, 0.05% Tween 20, pH 6.0) following staining with primary antibodies at 4°C overnight. Secondary antibodies were incubated for 60 min at room temperature (22–24°C). Slides were mounted with Faramount Aqueous Mounting Media (Cat. No. S3025, Dako). See supplemental for primary and secondary antibodies used.

### Morphological analysis

To quantify frequency and severity of pulmonary vascular changes in transgenic mice, whole sections of mouse lungs were stained with α-smooth muscle actin (Sigma-Aldrich, #2547) and von Willebrand factor (Dako, A0082) using standard immunohistochemical protocols. Scoring of peripheral muscularization based on 15 randomly acquired images per mouse. Peripheral vessels were defined as those small vessels distal to the terminal, muscularized bronchioles. Using a previously described method^41^, the degree of muscularization was defined by α-smooth muscle actin positivity around the vascular walls and classified as non-muscularized: < 20%, partial muscularization: 20-70%, fully muscularized: > 70%. In addition, sequential mouse lung specimens were stained with α-smooth muscle actin and von Willebrand factor with automated whole lung analysis of vasculatures performed by Visiomorph™ software (Visiopharm; Hoersholm, Denmark).

### Hemodynamic measurements (right heart catheterization)

Hemodynamic studies were performed by the Rodent Cardiovascular Phenotyping Core (RRID: SCR_022419) at the University of Pennsylvania. The mice were anesthetized with inhalation of isoflurane 3.5%, intubated, and mechanically ventilated (130 breaths per minute, FiO2 = 1.0). The right lateral neck was dissected to isolate the right internal jugular vein. A micromanometer pressure catheter (Millar SPR1000) was introduced into the right internal jugular vein and passed into the right ventricle for intraventricular arterial pressure monitoring. Right ventricular systolic pressure (RVSP), end diastolic pressure (EDP), heart rate, dP/dt were recorded and analyzed using a PowerLab 10 (AD Instruments, Colorado Springs, CO, USA). After data acquisition, the micromanometer pressure catheter was removed and the animals were euthanized.

### Measurement of RV hypertrophy (Fulton Index)

Hearts were excised and the RV free wall was dissected along the interventricular septum. RV hypertrophy was calculated as the weight ratio of the RV free wall relative to the left ventricle + septum (RV/LV + S). Measurements were captured using wet weights as well as dry weights with the latter obtained after specimens were dehydrated at 55C overnight.

### Isolation of primary ECs and fibroblasts

Single cell suspensions from distal human and whole mouse lungs were prepared using previously described methods.^21^ Immune cells were depleted using CD45-microbeads (Miltenyi, 130-045-801) and ECs were isolated after CD31+ selection with CD31+ microbeads (Milentyi, 130-091-935). ECs were grown on EGM2 (Lonza CC3162) on collagen-coated flasks. Fibroblast cells were derived from the CD45^-^ CD31^-^ fraction that were subsequently depleted of epithelial cells using EPCAM-conjugated Dynabeads (ThermoFisher 1120D). ECs were grown to confluence and sorted a second time with CD31+ microbeads prior to experimentation in passages 2-4.

### Endothelial-fibroblast coculture

Fibroblasts were plated on 12 well plates at 9×10^4^/well and incubated for 6 hours prior to the addition of ECs at 3×10^4^/well. Media (10% EBM + EGM2 without heparin) were changed every 2 days and the cocultures were grown for 6 days prior to fixation and staining.

### Transwell cell migration assay

Transwell inserts (8um pores; Corning 353097) were transferred into 24-well plates with culture medium. Cell suspension added to the upper chamber. Incubated at 37°C and 5% CO2 for 20 hours. Cells fixed with glutaldehyde and stained with crystal violet. Images taken on EVOS and cells/field counted.

### Proliferation assay

Primary cells between passages 2-4 were seeded in 96-well plates at 2.5 × 10^3^ cells/well in 200 μL of media with 2.5% FBS. Compounds were added the next day and further incubated for 48-versus 72-hours. Cells were fixed with 25% glutaraldehyde and stained with 0.05% crystal violet (C3886, Sigma-Aldrich, St. Louis, MO, USA). Cells were lysed with methanol prior to absorbance reading at 590nm.

### RNA isolation and qPCR analysis for gene expression

PureLink RNA Micro Kit (Invitrogen) was used to isolate RNA according to the manufacturer’s protocol. Determination of RNA concentration and purity was performed by optical density (OD) measurement (ratio of OD at 260 nm to OD at 280 nm is greater than 1.7) using a Nano-Drop spectrophotometer (Thermo Scientific). cDNA was synthesized using SuperScript IV Reverse Transcriptase (Thermo Fisher Scientific). qPCR was performed using 2× Power SYBR green reagents on the QuantStudio 7 Thermocycler (Life Technologies).

### Immunoblot analysis

Cells were washed with DPBS prior to lysis on ice for 15 mins in RIPA cell lysis buffer (Sigma) supplemented with protease and phosphatase inhibitors (Roche) as described [REF]. Samples were separated on SDS PAGE transferred to nitrocellulose, blocked and incubated overnight at 4 °C with the diluted primary antibody. Blots were washed then incubated for 60 min RT with the appropriate secondary antibody. Image acquisition and band intensity quantification was performed using an Odyssey IR imaging system (LiCor Biosciences Lincoln, NE).

### Statistics

Statistical analyses were performed in GraphPad Prism 5. A two-tailed Student’s *t*-test was used for the comparison between two experimental groups. For experiments with more than two groups, nonparametric Kruskal-Wallis ANOVA test was performed with Siegel (Bonferroni) correction for post-hoc, pair-wise contrasts. Statistical data were considered significant when *P* < 0.05.

## Acknowledgements

We thank the patients who contributed to this study, without which this study would not be possible. In addition, we thank the staff at the Rodent Cardiovascular Phenotyping Core for assistance with cardiac/hemodynamic measurements, the Pathology Core Laboratory at the Children’s Hospital of Philadelphia Research Institute and the Comparative Pathology Core at the University of Pennsylvania’s Veterinary School for providing histology services, National Disease Research Institute (NDRI, Philadelphia Pennsylvania) for providing human LAM lung explanted tissue specimens. Lastly, we would like to thank Dr. Slaven Crnkovic for his assistance with morphological analysis using the Visiomorph program.

This work was supported by grants from the National Institutes of Health including F32HL162425 (to S.M.L), 5T32HL007586-35 (to S.M.L, M.C.B, J.D.P), KL2TR001879 (to S.M.L and L.T.F), R03HL135227 (to E.C), R01HL151467, R01 HL158737, R01 HL141462, R41 HL156767, U01 HL131022 (to V.P.K). S.M.L was also supported by NIH LRP award.

## Contributions

SML designed and performed experiments, developed 2D and 3D ECs cocultures systems, performed IHC/IF, collected and measured Fulton Index, and wrote the manuscript. RR assisted with mouse care as well as mouse lung isolation. SML, MCB, LTF, JDP and EC provided access to human samples. AG and CS assisted with cell culture experiments. SC and GW performed Visiomorph analysis. KO processed mouse lung for bulk RNA sequencing. AM performed the RNAscope experiments. KSV performed the RT-PCR experiments. VPK designed the study, interpreted the data, wrote the manuscript, and directed the project. The authors would like to thank Dr. Michael Beers, Dr. Nilam Mangalmurti, Dr. Jacob Brenner and Dr. Andy Vaughn for helpful discussion of the data.

## Notes

### Competing Interest Statement

The authors have declared no competing interest.

