## Supplemental Figures for "Hyperactive mTOR in Lung Mesenchyme Induces Endothelial Dysfunction and Pulmonary Vascular Remodeling"

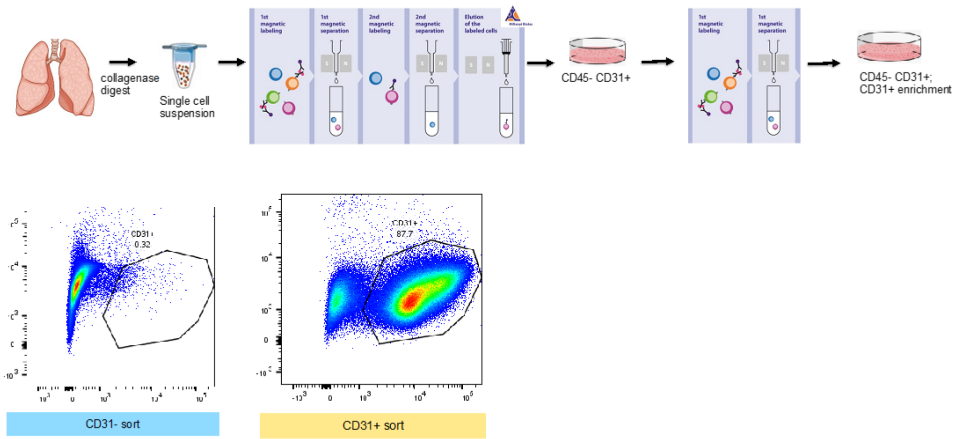


**Supplemental Figure 1. Isolation and purification of pulmonary ECs from lung explants of patients with LAM.** (a) Peripheral tissue are enzymatically digested and magnetically sorted using CD31+ microbeads. Primary cells are grown to confluence on T25 flask and then resorted using CD31+ microbeads. (b) Confirmation of purity of primary ECs isolates by flow cytometry. Flow cytometry of CD31+ sorted cells grown in primary cell culture with high enrichment (87.7%) versus CD31- depleted cells (0.32% positive for CD31).

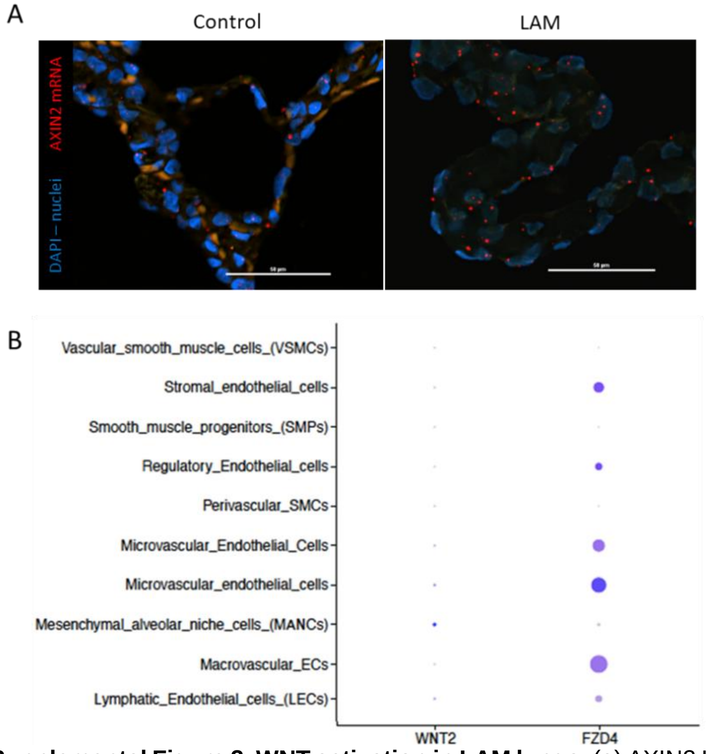


**Supplemental Figure 2. WNT activation in LAM lungs**. (a) AXIN2 In Situ Hybridization using RNA-scope in LAM and control human lung. AXIN2 probe (NM_015732, region 330–1287, Advanced Cell Diagnostics) was detected using Opal 570 and Opal 690 fluorophores (Akoya Biosciences). DAPI detects nuclei. (b) Wnt expression in mesenchymal and endothelial populations in control lung.

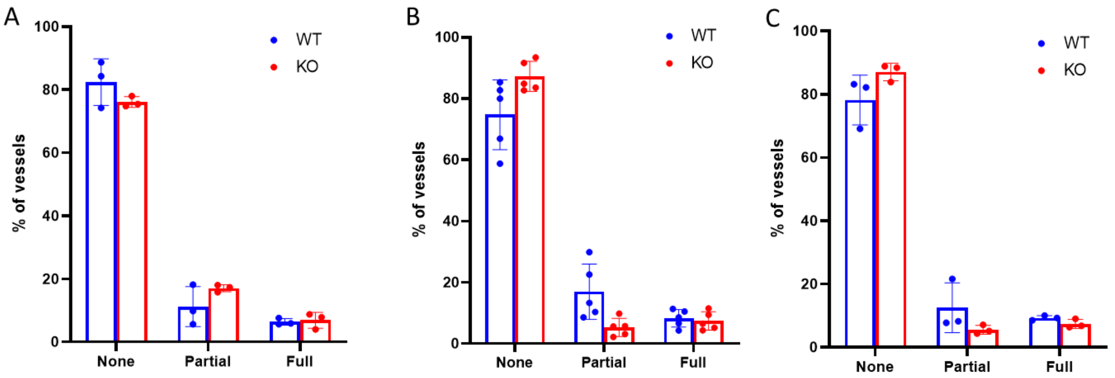


**Supplemental Figure 3**. **Vascular muscularization in young mouse lungs.** (a) 12-week-old, (b) 16-week-old and (c) 20-week old *Tsc2*$^{WT}$ and *Tsc2*$^{KO}$ mice. Muscularization was assessed using methods as previously described (n=3-5 animals/group). Statistical analysis were performed nonparametric Kruskal-Wallis ANOVA test with Siegel (Bonferroni) correction for post-hoc, pair-wise contrasts.

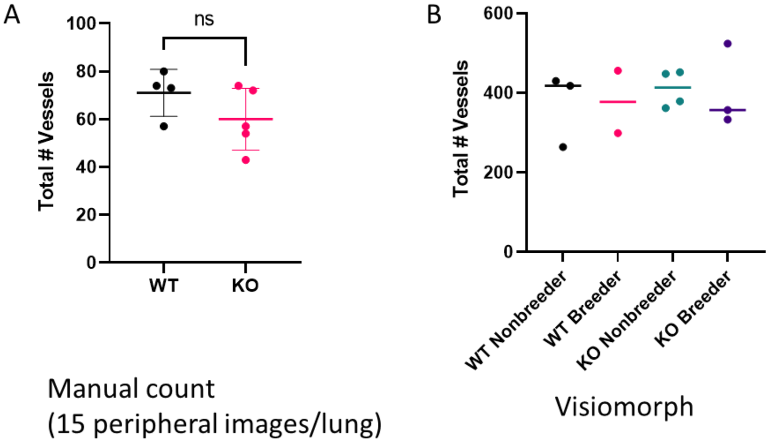


**Supplemental Figure 4. Total vessels in one-year-old mice** as assessed by (a) manual counts (n=4 *Tsc2*$^{WT}$, 5 *Tsc2*$^{KO}$) and (b) Visiomorph analysis (n=5 *Tsc2*$^{WT}$, 7 *Tsc2*$^{KO}$). Statistical analysis was performed nonparametric Kruskal-Wallis ANOVA test.

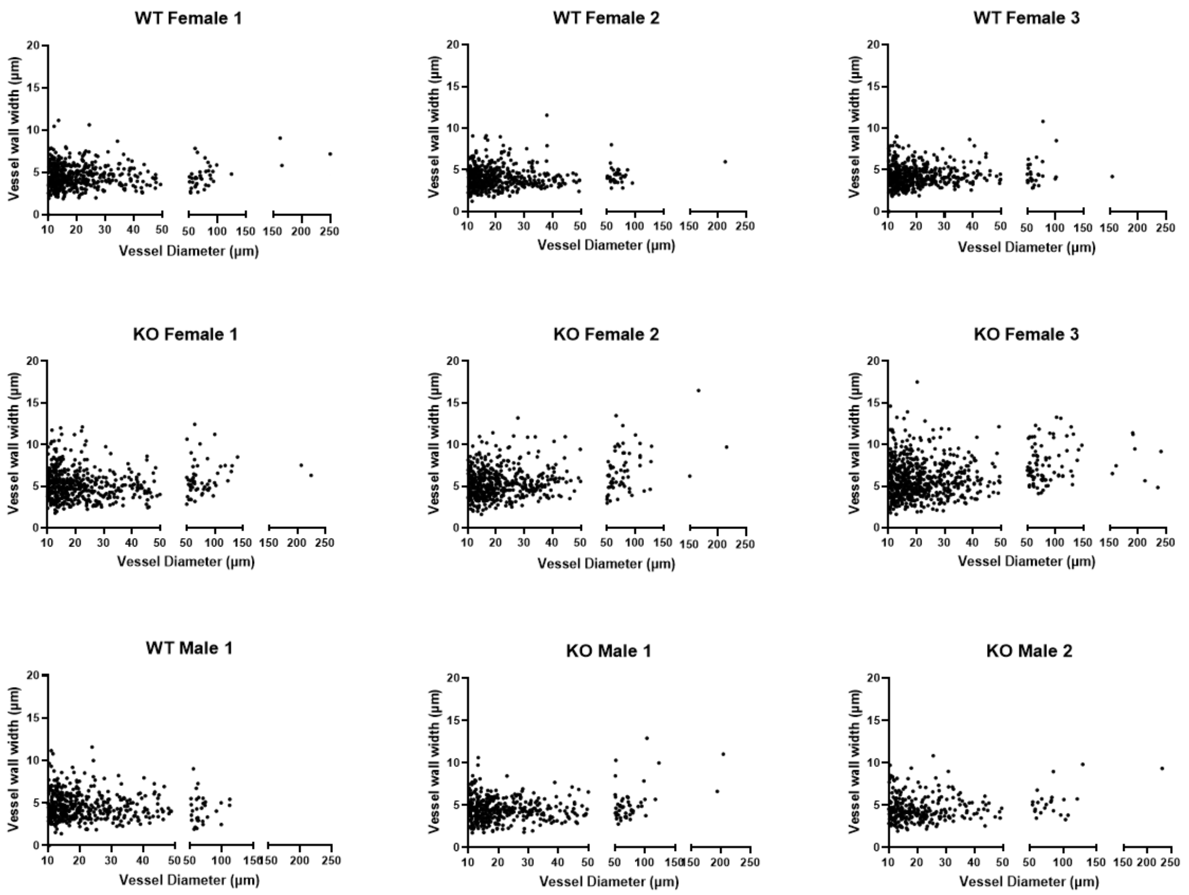


**Supplemental Figure 5. Distribution of vessel wall thickness individual mouse lungs.** Vessel thickness were calculated as the distance between border of the vessel wall of the lumen to the vessel wall-lung tissue interface (encompassing the medial and intimal layer). *Tsc2*$^{KO}$ mice had thicker vessels compared to age matched controls.

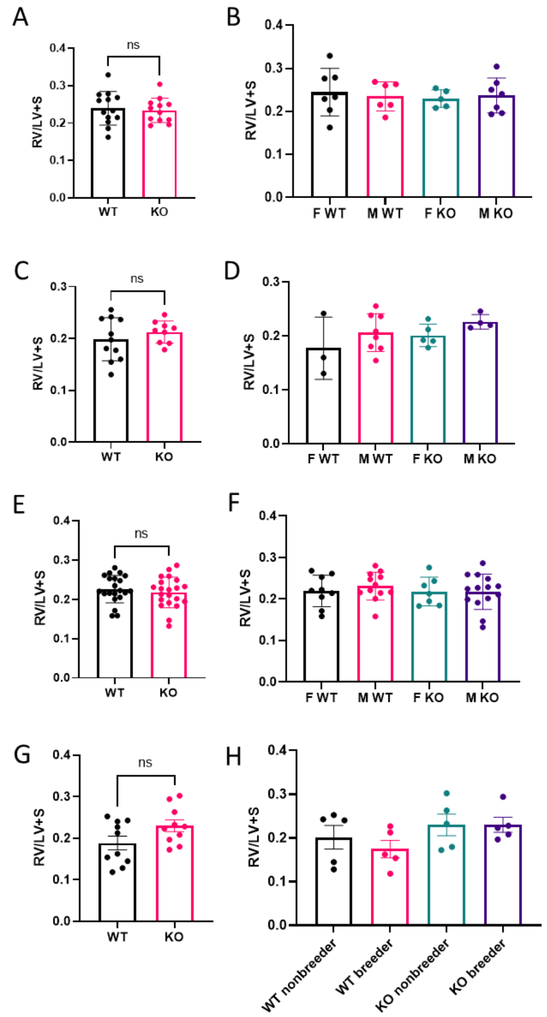


**Supplemental Figure 6. Fulton Index in 12-, 16- and 24-week old *Tsc2^WT^* and *Tsc2^KO^* mice.** Fulton Index was assessed as described in Methods. (a) 12-week-old mice, n= 13 WT, 12 K). (b) 12-week-old mice by gender, n= 7 F WT, 5 M WT, 5 F KO, 7 M KO. (c) 16-week-old mice, n= 11 WT, 9 KO. (d) 16-week-old by gender, n=3 F WT, 8 M WT, 5 F KO, 4 M KO. (e) 20-week old mice, n= 21 WT, 21 KO. (f) 20-week-old by gender, n= 9 F WT, 12 M WT, 7 F KO, 14 M KO. (g) 24-week-old female mice, n = 10 WT, 10 KO. (h) 24-week-old breeder mice, n= 5 F WT, 5 M WT, 5 F KO, 5 M KO. Statistical analysis was performed using two-tailed Student's *t*-test or nonparametric Kruskal-Wallis ANOVA test.

**RT-PCR primers:**

| Gene | Forward Primer sequence 5'-->3' | Reverse Primer sequence 5'-->3' |
|---|---|---|
| Axin2 (mouse) | ATTTTCCGAGAACCCACCGC | TCCTTTTCTTCATCCTCCCGG |
| LRP6 (mouse) | AGTCAGTTTGTGGTCACGGC | GTCCAATACATGTACCCAACCA |
| Fzd1 (mouse) | TGTACTTCGGGCCAGAGGAG | CGGTCCTCCAACAGAAAGCC |
| Fzd2 (mouse) | CTCTGGGCAGGCTCTTTGTG | AGAACATCTCTCAGGCCGCT |
| Fzd7 (mouse) | GACCAAGCCATTCCTCCGTG | GGGGACATCGCAGTGAAAGG |
| Fzd8 (mouse) | AGATACAGTGCTCCCCGGAC | GCGGTTGTAGTCCATGCACA |
